# Engineered Th17 cell differentiation using a photo-activatable immune modulator

**DOI:** 10.1101/719021

**Authors:** Bibudha Parasar, Pamela V. Chang

## Abstract

T helper 17 (Th17) cells, an important subset of CD4^+^ T cells, help to eliminate extracellular infectious pathogens that have invaded our tissues. Despite the critical roles of Th17 cells in immunity, how the immune system regulates the production and maintenance of this cell type remains poorly understood. In particular, the plasticity of these cells, or their dynamic ability to trans-differentiate into other CD4^+^ T cell subsets, remains mostly uncharacterized. Here, we report a synthetic immunology approach using a photo-activatable immune modulator (PIM) to increase Th17 cell differentiation on demand with spatial and temporal precision to help elucidate this important and dynamic process. In this chemical strategy, we developed a latent agonist that, upon photochemical activation, releases a small-molecule ligand that targets the aryl hydrocarbon receptor (AhR) and ultimately induces Th17 cell differentiation. We used this chemical tool to control AhR activation with spatiotemporal precision within cells and to modulate Th17 cell differentiation on demand by using UV light illumination. We envision that this approach will enable an understanding of the dynamic functions and behaviors of Th17 cells in vivo during immune responses and in mouse models of inflammatory disease.

## Introduction

Synthetic immunology harnesses the ability to manipulate immune cells using rational design of synthetic systems to either understand their functions or treat inflammatory diseases^1–4^. The human immune system represents an incredibly complex consortium of cell types that orchestrates immune responses to foreign invaders, including pathogenic microbes. Upon sensing these microorganisms, the host immune system initiates many signaling pathways to activate different immune cell types to help clear the infection^5,6^. This multifaceted process involves both spatial and temporal positioning of a network of cells to facilitate information exchange. Such cell-cell interactions ensure an effective response because the various cell types communicate with one another through the expression of key molecules that are either secreted or displayed on their cell surfaces. Due to the sheer number of cells and cell types that are involved in the immune response, understanding the precise roles of each cell type in this highly dynamic and interactive process is a challenging task. Thus, new strategies are needed to control the host immune response to help elucidate the functions of various immune cells and the inflammatory pathways that they activate in vivo.

T helper 17 (Th17) cells are a subset of CD4^+^ T cells whose main roles are to assist in the eradication of extracellular pathogens that are typically associated with mucosal surfaces^7^. When left unchecked by the immune system, these pro-inflammatory CD4^+^ T helper cells can contribute to allergy and autoimmune diseases due to increased production of pro-inflammatory cytokines such as IL-17. The differentiation of naïve T cells into Th17 cells depends on the presence of specific cytokines secreted by activated antigen-presenting cells, including IL-6 and TGF-β, and key transcription factors that define the Th17 lineage are the retinoic acid receptor-related orphan receptors (ROR)-gamma (RORγ) and RORα^8,9^.

The aryl hydrocarbon receptor (AhR) is a ligand-activated transcription factor whose classic roles include activation of the host xenobiotic response to eliminate noxious chemicals from the body^10^. The main target genes of AhR include those that encode cytochrome P450 enzymes, which oxidize these foreign compounds into metabolites that can be more easily excreted by the host^11^. In recent years, immunologists have discovered that AhR also plays a major role in the regulation of numerous immune cell types, including the differentiation of specific CD4^+^ T cell subsets such as Th17 cells^12,13^. In particular, AhR can enhance Th17 cell differentiation via its activation by specific ligands derived from tryptophan, including 6-formylindolo[3,2-b]carbazole (FICZ), and chronic AhR activation controls the trans-differentiation of Th17 cells into anti-inflammatory T regulatory type 1 cells, which suppress pro-inflammatory T cell responses in vivo^14,15^.

Despite these recent advances, understanding Th17 cell plasticity, i.e., their ability to trans-differentiate into distinct T cell subsets in vivo, has been hindered by a lack of technologies to interrogate or manipulate their activities in vivo under physiologically relevant conditions^16,17^. This need has been partially met by fate mapping using fluorescent reporter mice; however, these genetic approaches suffer from limited spatial and temporal resolution^18^. Here, we have developed a chemical strategy suitable for use in vivo for the precise control of Th17 cell differentiation via activation of AhR. In this approach, we harness the exquisite control afforded by light irradiation to selectively release the AhR ligand FICZ via photochemistry with spatiotemporal precision.

Synthetic immunology approaches exploiting photo-activation have been utilized to modulate additional immune cell types, including dendritic cells and macrophages, by targeting different signaling pathways^19–22^. Alternatively, small-molecule approaches using bioorthogonal chemical deprotection have also been developed to activate cytotoxic CD8^+^ T cells, a different T cell type^23,24^. However, there exist no reports of chemical strategies to manipulate CD4^+^ Th17 cells or the AhR pathway with dynamic control. We envision that our strategy to selectively induce Th17 cell differentiation on demand will help elucidate a mechanistic understanding of Th17 cell plasticity in vivo by probing Th17 cell activity during immune responses to pathogens and in mouse inflammatory disease models.

## Results and Discussion

Toward this ultimate goal, we have developed a **P**hoto-activatable **I**mmune **M**odulator of **Th17** cells (PIM-Th17, Figure 1), which releases FICZ upon UV light illumination. To control the release of FICZ in a spatiotemporal manner, we modified the ligand with a well-characterized photo-protecting group, 6-nitroveratryl (NV)^25^, to render it inactive to AhR. Upon mild light illumination (365 nm), the NV group is cleaved by photolysis and FICZ is released, allowing its selective delivery to the target cells. We synthesized PIM-Th17 from FICZ in a one-step displacement reaction using 6-NV-bromide (Scheme S1).

**Figure 1.**
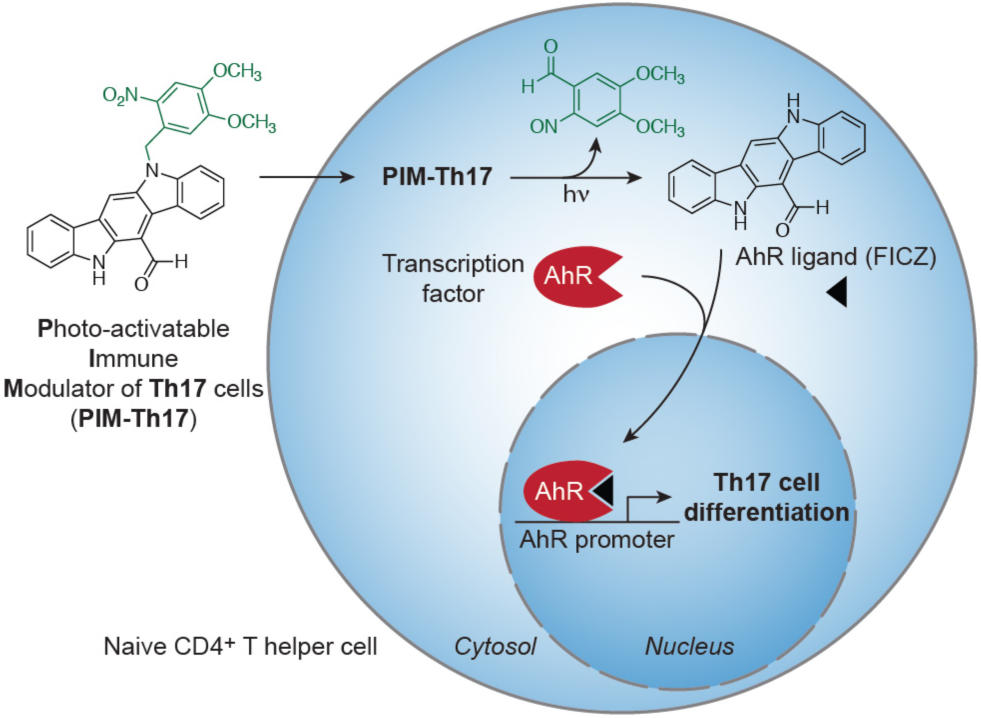
Schematic of synthetic immunology approach for inducing T helper 17 (Th17) cell differentiation by activation of the aryl hydrocarbon receptor (AhR) pathway. UV light illumination of a photocaged AhR ligand, **P**hoto-activatable **I**mmune **M**odulator of **Th17** cells (PIM-Th17), releases the AhR ligand, FICZ, which binds AhR. Ligand binding activates nuclear translocation and induction of Th17 cell differentiation.

We first demonstrated that PIM-Th17 can be photo-uncaged using UV light. In these studies, we illuminated PIM-Th17 with UV light (365 nm) and monitored its conversion to FICZ over time using HPLC, followed by mass spectrometry (Figure 2). We found that photo-uncaging of PIM-Th17 followed first order reaction kinetics, as expected, with a rate constant of 4.85×10^-3^ s^-1^ (Figure S1). Although the major product formed was FICZ, we also observed the expected nitrosoaldehyde and several minor byproducts (Figure S2A)^25^. We determined that none of these byproducts activated AhR using a luciferase reporter assay for AhR activity (Figure S2B)^26^. We also demonstrated that PIM-Th17 does not activate AHR until it is illuminated with UV light, which releases FICZ (Figure S3). Furthermore, PIM-Th17 is stable to aqueous hydrolysis for at least 4 days at 37 °C (Figure S4).

**Figure 2.**
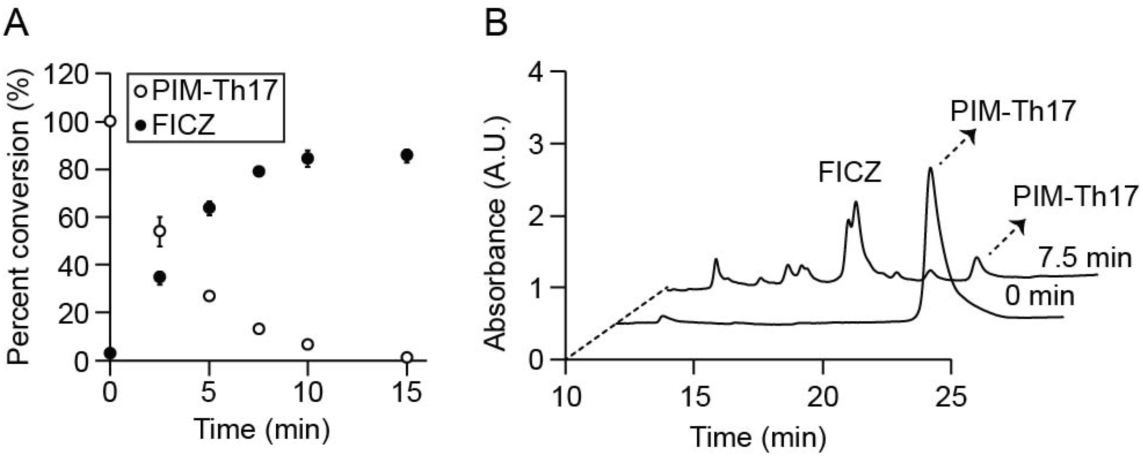
PIM-Th17 is photo-uncaged by UV light. (A) Percent conversion of PIM-Th17 to FICZ after illumination with 365 nm light. (B) HPLC analysis of photo-uncaging reactions. Traces shown were monitored by UV absorbance at 254 nm. A.U. = arbitrary units.

Next, we determined whether photo-uncaging of PIM-Th17 leads to AhR nuclear translocation in mammalian cells (Figures 3, S5)^27^. We transfected HEK 293T cells with both a yellow fluorescent protein fusion to AhR (AhR-YFP) and photo-activatable mCherry (PA-mCherry), a fluorogenic protein that enables tracking of cells that were illuminated with UV light^28^. We then incubated the reporter cells with PIM-Th17 for 30 min to allow for cellular uptake, followed by a quick wash and illumination of a population of cells within the center of the dish (Figure 3A). Using confocal microscopy, we found that photo-uncaging of PIM-Th17 led to increased AhR nuclear translocation in the cells that had been illuminated in the center of dish compared to those in the periphery that had not been illuminated (Figure 3B-C). To demonstrate the precise spatial selectivity of this approach, we also imaged adjacent regions of the dish that had been illuminated and non-illuminated (Figure 3D, dashed line denotes border region of illumination). Importantly, neither UV irradiation nor the concentrations used of PIM-Th17 were toxic to the cells as determined by propidium iodide staining (Figure S6A).

**Figure 3.**
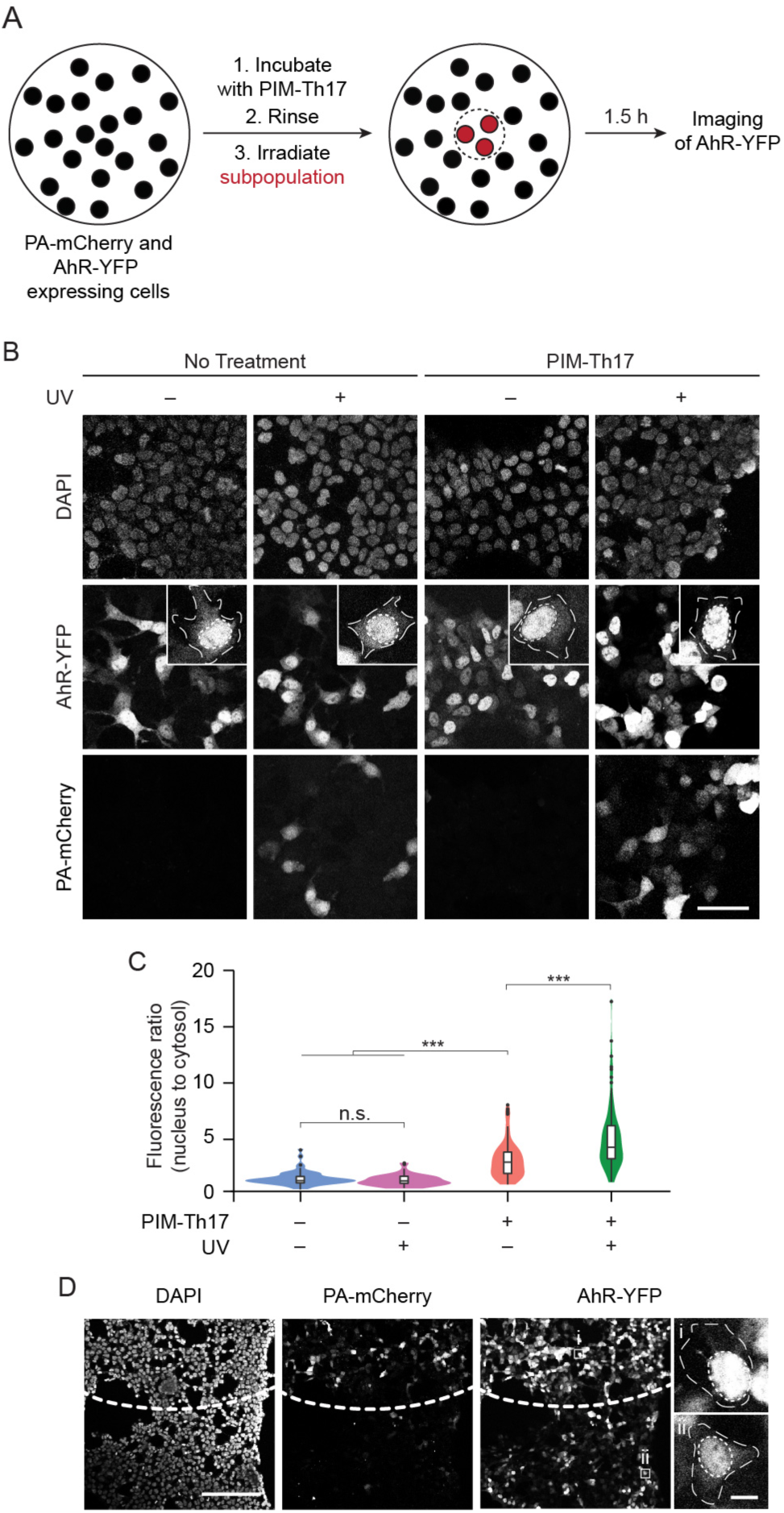
Photo-uncaging of PIM-Th17 is spatially selective and leads to nuclear translocation of AhR. HEK 293T cells co-transfected with AhR-YFP and PA-mCherry were pretreated with PIM-Th17 (10 µM) or vehicle (DMSO) for 30 min, and the center of the dish was illuminated with UV light (epifluorescence using DAPI filter, 30 s). Cells were subsequently incubated for an additional 1.5 h, fixed, stained with DAPI, and imaged. (A) Schematic of experimental setup. (B) Maximum intensity z-projection images of cells at the periphery (– UV) or center (+ UV) of the same dish, imaged by confocal microscopy. Representative individual cells are shown at higher magnification (3.85X) in inset images. (C) Violin plots of nuclear to cytosolic ratio of the AhR-YFP fluorescence from (B), n = 50–70 cells in each treatment group. Box plot inside the violin plot: Interquartile ranges (IQRs, boxes), median values (line within box), whiskers (lowest and highest values within 1.5 times IQR from the first and third quartiles), and outliers beyond whiskers (dots) are indicated. Statistical significance was assessed using one-way ANOVA followed by post-hoc Tukey’s test. n.s. = not significant, \*\*\**p* < 0.001. (D) Maximum intensity z-projection images of cells at the border of UV illumination. Dashed curve indicates border region of illumination (+ UV, top; – UV, bottom), and representative individual cells are indicated by white squares and shown at higher magnification at right. Scale bars: 50 µm (B and D, full-size images), 10 µm (D, magnified images).

We then determined that photo-uncaging of PIM-Th17 leads to increased transcription of canonical AhR target genes, including those that encode cytochrome P450 enzymes and the aryl hydrocarbon receptor repressor (AhRR)^11^. In these studies, HepG2 cells were treated with and without PIM-Th17, followed by UV light illumination. We found that photo-uncaging of PIM-Th17 leads to increased transcript levels of *Cyp1a1, Cyp1a2, Cyp1b1*, and *Ahrr* by qPCR (Figure 4). Collectively, these results suggest that AhR nuclear translocation caused by photo-uncaged PIM-Th17 results in binding of AhR to its transcription factor binding site and increased transcription of downstream target genes. We also verified that the concentrations of PIM-Th17 and UV illumination used in this experiment were not toxic to HepG2 cells (Figure S6B).

**Figure 4.**
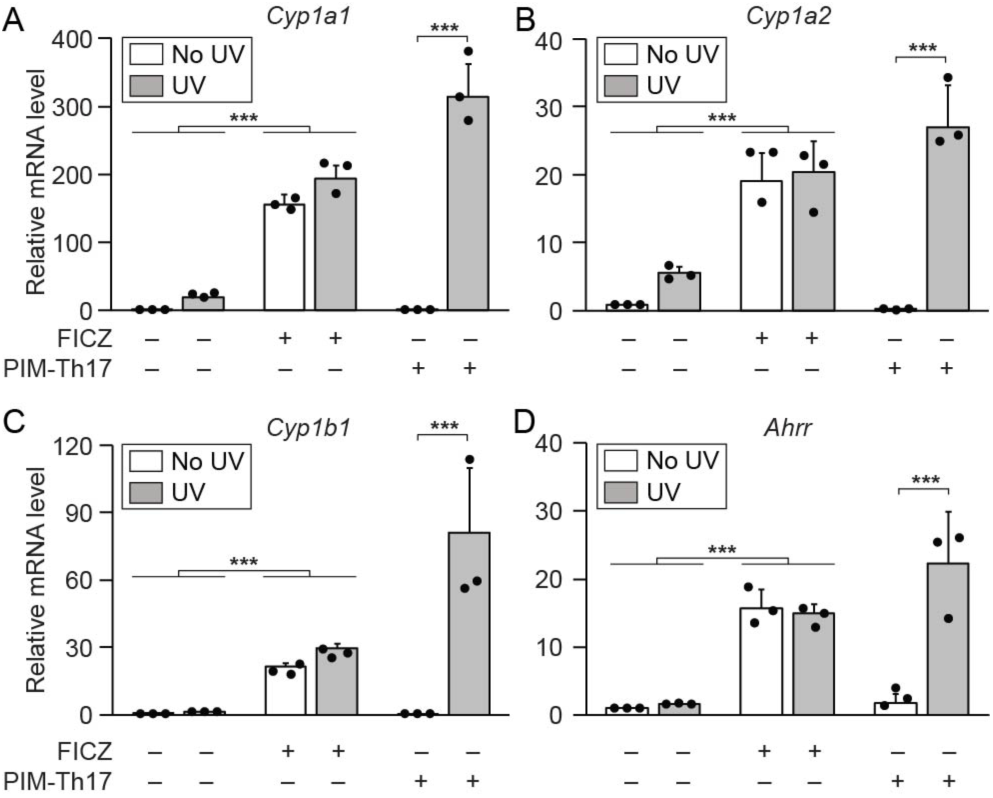
Photo-uncaging of PIM-Th17 leads to transcription of AhR downstream target genes. HepG2 cells were treated with FICZ (300 nM), PIM-Th17 (10 µM), or vehicle (DMSO) and illuminated with UV light (365 nm, 10 min, grey bars) or kept in the dark (white bars). RNA was isolated after 8 h, and cDNA was synthesized and analyzed by qPCR (data normalized to *Rpl13a*). Relative mRNA transcript levels of (A) *Cyp1a1*, (B) *Cyp1a2*, (C) *Cyp1b1*, and (D) *Ahrr* are shown. Statistical significance was assessed using one-way ANOVA followed by post-hoc Tukey’s test. \*\*\**p* < 0.001.

Finally, we evaluated the ability of PIM-Th17 to increase differentiation of Th17 cells upon photo-uncaging. We isolated naïve CD4^+^ T cells from the spleens of wild-type mice and cultured these cells under Th17 differentiation conditions in the presence or absence of PIM-Th17^12^. Importantly, we verified that the concentrations of PIM-Th17 and UV illumination used were not toxic to CD4^+^ T cells (Figure S6C). Gratifyingly, after photo-uncaging of PIM-Th17 to release FICZ, we found that the percentage of Th17 cells within the T cell population increased compared to that in a parallel population of T cells that were treated with PIM-Th17 but were not illuminated (Figure 5). These results demonstrate that our PIM-based chemical strategy can be used to selectively differentiate Th17 cells on demand via activation of AhR.

**Figure 5.**
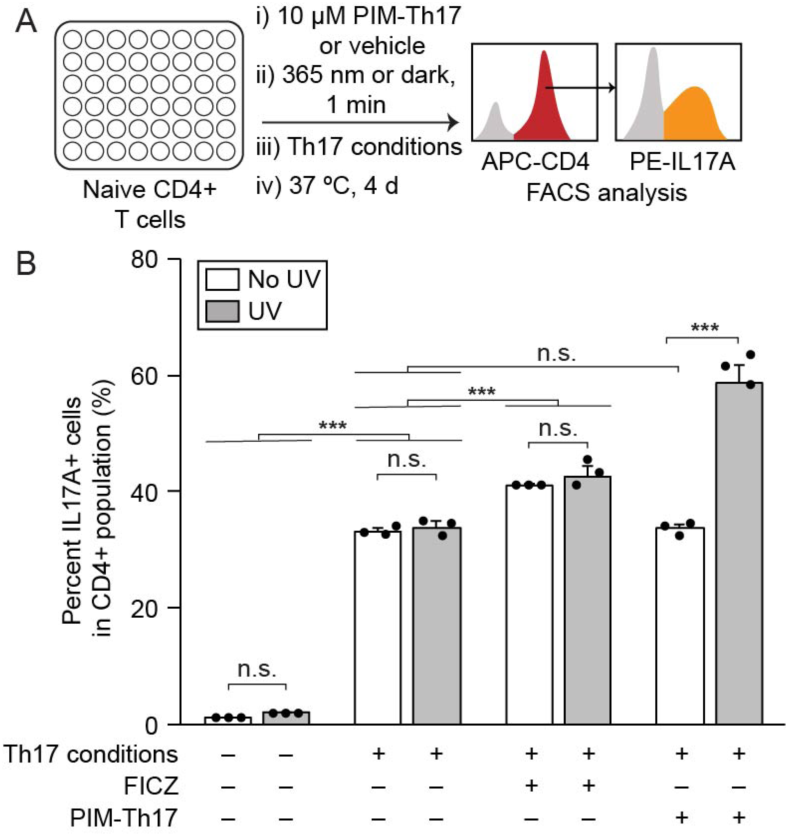
Photo-uncaging of PIM-Th17 increases differentiation of Th17 cells from naïve CD4^+^ T cells. Splenic naïve CD4^+^ T cells were harvested from wildtype mice and cultured with or without Th17-inducing factors (Th17 conditions) in the presence of PIM-Th17 (10 µM), FICZ (300 nM), or vehicle (DMSO). The cells were illuminated with UV light (365 nm, 1 min) or kept in the dark, and then incubated at 37 °C for 4 d. On day 4, cells were stained with allophycocyanin (APC) anti-CD4 and phycoerythrin (PE) anti-IL17A and analyzed by FACS. (A) Schematic of experimental setup. (B) Quantification of percent CD4^+^IL17^+^ cells by FACS analysis. Statistical significance was assessed using one-way ANOVA followed by post-hoc Tukey’s test. n.s. = not significant, \*\*\**p* < 0.001.

## Conclusions

We have developed a synthetic immunology approach that utilizes a photo-activatable immune modulator (PIM) to control the differentiation of Th17 cells by targeting the AhR pathway with spatial and temporal precision. This chemical strategy exploits the use of photochemistry to selectively deliver an AhR ligand to the target cell type, which will enable manipulation of CD4^+^ T cell populations on demand to understand their dynamic roles in vivo. Additionally, this approach can be used to activate AhR within alternative cell types, including those involved in xenobiotic metabolism and additional immune cells that are regulated by this pathway^10^. Ultimately, we envision that this technology can be used to control Th17 cells during active immune responses in vivo and in mouse models of inflammatory disease to better understand the plasticity of this enigmatic immune cell type during various physiological and pathological states.

## Acknowledgments

This work was supported by the Arnold and Mabel Beckman Foundation (Beckman Young Investigator Award to P.V.C.) and a President’s Council of Cornell Women Affinito-Stewart Grant (P.V.C.). We thank the Cornell University NMR facility (NSF MRI: CHE-1531632) and Dr. Ivan Keresztes for technical assistance. We are grateful to Samantha Scott for animal husbandry and Dr. Brittany White for helpful discussions. We also thank the Weill Institute for Cell and Molecular Biology for resources and reagents.

