## Supporting Information for "Engineered Th17 cell differentiation using a photo-activatable immune modulator"

**Contents**

p. 2 **Materials and methods**
p. 6 **Synthetic procedures**
p. 7 **Figure S1.** Photo-uncaging of PIM-Th17 is a first-order reaction
p. 8 **Figure S2.** Photo-uncaging of PIM-Th17 produces minor byproducts that do not activate AhR
p. 9 **Figure S3.** PIM-Th17 does not increase AhR activity until photo-uncaged
p. 9 **Figure S4.** PIM-Th17 is stable in aqueous solution
p. 10 **Figure S5.** FICZ leads to nuclear translocation of AhR-YFP
p. 11 **Figure S6.** Photo-uncaging conditions are not toxic to cells
p. 12 **Figure S7:** $^{1}H$ NMR spectrum of PIM-Th17
p. 13 **Figure S8:** $^{13}C$ NMR spectrum of PIM-Th17
p. 14 **Figure S9:** 2D ROESY spectrum of PIM-Th17
p. 15 **Supplemental references**

### Materials and methods

**General materials and methods.** All chemical reagents were of analytical grade, obtained from commercial suppliers, and used without further purification unless otherwise noted. Organic extracts were dried over $Na_2SO_4$, and solvents were removed with a rotary evaporator at reduced pressure (20 torr), unless otherwise noted. Flash chromatography was performed using Silicycle Siliaflash P60 40-63 Å 230-400 mesh silica gel. Analytical thin layer chromatography (TLC) was performed on glass-backed TLC 60 Å silica gel plates, and compounds were visualized by staining with ceric ammonium molybdate and the absorbance of UV light (λ = 254 nm or 365 nm). Reverse phase HPLC was performed using a Shimadzu system equipped with a CBM-20A controller, SPD20AV UV-Vis detector, LC-20AR liquid chromatograph unit, FRC-10A fraction collector, and an Epic Polar 5 µm 120 Å C18 analytical column (4.6 x 250 mm) at a flow rate of 1 mL/min or a semipreparative column (10 x 250 mm) at a flow rate of 5 mL/min. HPLC samples were filtered with a Millex-LH syringe filter equipped with a 0.45 µm PTFE membrane prior to injection. The dd$H_2O$:acetonitrile gradient varied from 80:20 to 35:65 over 25 min with a curve value of -6, after which the gradient increased to 0:100 over 2 min at a linear rate and remained at 0:100 for an additional 4 min. 4,5-Dimethoxy-2-nitrobenzyl bromide (NV-Br) was purchased from Acros Organics. Sodium hydride (60% dispersion in mineral oil) was purchased from TCI America. Ethyl acetate, hexane, dimethylsulfoxide (DMSO), dimethylformamide (DMF), HPLC-grade dd$H_2O$ and acetonitrile were purchased from Fisher Scientific. NMR spectra were collected on a Varian INOVA 500 or 600 spectrometer and are referenced to residual solvent peaks. High resolution mass spectrometry (HRMS) was performed on an Agilent 6230 electrospray ionization–time-of-flight (ESI-TOF) mass spectrometer.

Dulbecco's phosphate-buffered saline (PBS) pH 7.4, RPMI-1640 media, L-glutamine, sodium pyruvate, HEPES buffer, and penicillin/streptomycin were purchased from Corning. Fetal bovine serum, Lipofectamine 2000, and ProLong Diamond mounting media with DAPI were purchased from Thermo Fisher Scientific. Tissue culture-treated dishes and cell strainers (70 µm) were obtained from BD Falcon. Glass-bottom microwell dishes (35 mm) were obtained from MatTek Corporation. Plasmids pGudluc6.1 and pRLTK were obtained from W. Eddie Ip (Mayo Clinic), and pAHR-YFP was obtained from Gary Perdew (Pennsylvania State University). pPAmCherry1-N1 was purchased from Addgene (plasmid #31928). ACK lysis buffer was obtained from Lonza. Naïve mouse $CD4^+$ T cell isolation kit, MACS multi-stand with an octo-MACS separator magnet, and MACS separation columns (high gradient, type MS) were purchased from Miltenyi Biotec. Recombinant mIL-6 and hTGF-β were purchased from R&D Systems. Rat anti-mouse IL-4 (NA/LE, clone 11B11) and rat anti-mouse IFN-γ (NA/LE, clone R4-6A2) were purchased from BD Biosciences. Anti-mouse CD28 (clone 37.51) and anti-mouse CD3e (clone 145-2C11) were purchased from BioLegend. Anti-mouse CD16/CD32 (clone 93), APC anti-mouse CD4 (clone GK1.5), PE anti-mouse/rat IL-17A (clone eBio17B7), IC fixation and permeabilization buffer set, and propidium iodide were purchased from Affymetrix eBioscience. Brefeldin A, phorbol 12-myristate 13-acetate, and ionomycin were purchased from Cayman Chemical. RNA-Bee was purchased from Tel-Test. PerfeCta SYBR Green SuperMix, Low ROX was obtained from Quanta Biosciences. SMART MMLV reverse transcriptase was purchased from Clontech. Glycogen (from mussels) was obtained from Roche. Imaging was performed on a Zeiss LSM 800 confocal laser scanning microscope equipped with 20X 0.8 NA and 40X 1.4 NA Plan Apochromat objectives, 405, 488, 561, and 640 nm solid-state lasers, and two GaAsP PMT detectors. Image analysis was performed using the Zeiss Zen Blue 2.3 and

FIJI software packages. Images shown are maximum intensity z-projection images, and quantification was performed using FIJI. Flow cytometry was performed on a BD FACS Canto II. Luminescence was collected on a Veritas microplate luminometer (Turner Biosystems). Samples were irradiated with a UVP Transilluminator 2UV.

**Mice.** C57Bl/6 mice were purchased from Jackson Laboratories and bred at the animal facility of Cornell University. Mice were used at 8-12 weeks of age in accordance with the guidelines of the Institutional Animal Care and Use Committee and the Cornell Center for Animal Resources and Education (Protocol number 2015-0069).

**In vitro photo-uncaging of PIM-Th17.** PIM-Th17 was dissolved in DMSO and irradiated with 365 nm UV light for 15 min. The solution was periodically analyzed by HPLC and MS to monitor conversion of PIM-Th17 to FICZ. Percent conversion was calculated by analyzing standards of PIM-Th17 and FICZ by HPLC, using integration of the area under the curve for normalization. These data were used to calculate the first-order reaction kinetics.

**In vitro stability studies.** PIM-Th17 was dissolved in a mixture of 5:1 DMSO:PBS and incubated at 37 °C for 4 d. The solution was periodically analyzed by HPLC and MS to monitor decomposition of PIM-Th17. Percent of PIM-Th17 remaining was calculated by analyzing a standard of PIM-Th17 by HPLC, using integration of the area under the curve for normalization.

**Photo-uncaging of PIM-Th17 and identification of byproducts by mass spectrometry.** PIM-Th17 (4.4 mM in DMSO) was irradiated with UV light (365 nm, 10 min). The reaction products were purified by HPLC and analyzed by HRMS to identify the byproducts.

**AhR luciferase reporter assay.** Passive lysis buffer (5x PLB) was prepared with 125 mM Tris, pH 7.8, 10 mM 1,2-diaminocyclohexane tetraacetic acid (CDTA), 10 mM DTT, 5 mg/mL BSA, 5% (vol/vol) Triton X-100, and 50% (vol/vol) glycerol in $ddH_2O$. An aqueous solution of 1x firefly luciferase substrate was prepared containing 75 mM HEPES, pH 8.0, 4 mM $MgSO_4$, 20 mM DTT, 0.1 mM EDTA, 0.53 mM ATP, 0.27 mM coenzyme A, and 0.47 mM D-luciferin (firefly) in $ddH_2O$. An aqueous solution of 1x Renilla luciferase buffer was prepared containing 7.5 mM sodium acetate, pH 5.0, 400 mM sodium sulfate, 10 mM CDTA, 15 mM sodium pyrophosphate, and 0.025 mM 2-(4-aminophenyl)-6-methylbenzothiazole. A 100x Renilla luciferase substrate was prepared by diluting coelenterazine to 0.55 mM in anhydrous methanol and added to 1x Renilla luciferase buffer immediately prior to the assay.

PIM-Th17 (10 μM) was photo-uncaged in vitro at 37 °C for 3 h, and the byproducts were purified by HPLC. HEK 293T cells were plated ($2x10^5$ cells/well) in a 24-well plate. After the cells reached 50-60% confluency, they were transfected with 1 μg of pGudluc6.1[1] and 100 ng of pRLTK per well. Following transfection, the cells were treated with FICZ (300 nM), PIM-Th17 (10 μM) with and without UV light (365 nm, 10 min), or the isolated byproducts at concentrations based on photo-uncaging of PIM-Th17 as above for 3 h. Cells were washed with 1X PBS and lysed with 250 μL of 1x PLB. Twenty microliters of the lysate were added to a 96-well white opaque plate (Corning). After adding 50 μL of 1x firefly luciferase substrate, luminescence was measured for 2 min. Subsequently, 50 μL of the 1x Renilla substrate was added to each sample, and luminescence was measured after 2 min. The luciferase activity was calculated by calculating the ratio of firefly luciferase signal to Renilla luciferase signal.

**Cell viability assay.** For HeLa, HEK 293T, and HepG2 cells: Cells were plated in a 96-well plate with a density of $2.5x10^4$ cells/well. After 24 h, cells were either incubated with various concentrations of PIM-Th17 or irradiated with or without UV light (365 nm) for different periods of time. After the appropriate treatment, cells were incubated at 37 °C for 24 h prior to the assay.

For T cells: Splenic naïve $CD4^+$ T cells (isolated as described below) were plated in a 96-well plate with a density of $2.5x10^5$ cells/well. Cells were then either incubated with various concentrations of PIM-Th17 or irradiated with or without UV light (365 nm) for different periods of time. After the appropriate treatment, cells were incubated at 37 °C for 4 d prior to the assay.

Propidium iodide (PI) staining: After incubation, cells were collected into microcentrifuge tubes and transferred to a 96-well V bottom plate and washed with 170 µL of 1X PBS. Cells were pelleted at 350 ×g for 3 min at 4 °C. PI was diluted (1:40) in FACS buffer (1X PBS + 0.5% FBS), and 50 µL of the PI working reagent was added to each well. Cells were gently mixed by pipetting up and down and were transferred to a FACS tube containing 150 µL of FACS buffer. Cells were then analyzed by FACS, and percent live cells was calculated.

**RNA purification and qPCR analysis.** HepG2 cells were seeded ($2.5x10^5$ cells/well) in a 24-well plate. After cells were confluent, they were treated in the presence of FICZ (300 nM), PIM-Th17 (10 µM), or vehicle (DMSO). Cells were incubated at 37 °C for 8 h after which total RNA was isolated with RNA-Bee reagent according to the manufacturer's protocol. Total RNA was reverse transcribed with an oligo(dT) primer using MMLV RT. cDNA was analyzed in triplicate by qPCR amplification using SYBR Green Supermix on a Bio-Rad CFX96 Real-Time PCR Detection System. PCR amplification conditions were as follows: 95 °C (3 min) and 40 cycles of 95 °C (15 s) and 60 °C (45 s). Primer pairs (for sequences see Table S1) were designed to amplify mRNA-specific fragments, and unique products were tested by melt-curve analysis [95 °C (15 s), 60–95 °C (Δ 0.5 °C, 5 s)]. Relative expression was normalized to ribosomal protein L13a (*Rpl13a*). Data are represented as the fold induction over untreated cells.

**Photo-uncaging of PIM-Th17 and confocal microscopy of AhR-YFP reporter cells.** HEK 293T cells were plated ($2x10^5$ cells/well) in 35 mm glass-bottom microwell dishes. After the cells reached 50-60% confluency, each well was transfected with 1.5 µg of pAhR-YFP[2] and 1.5 µg of pPAmCherry1-N1[3]. The cells were then incubated for 12 h after which the cells were treated with FICZ (300 nM), PIM-Th17 (10 µM), or vehicle (DMSO) for 30 min. Cells were subsequently washed with PBS (1 x 2 mL), and the PBS was replaced with 2 mL of DMEM (with 10% FBS and penicillin/streptomycin). The cells were irradiated with UV light using epifluorescence illumination (HXP 120 V light source) with a 10X objective at 100% intensity through a DAPI filter set (Zeiss #49) for 30 s. Cells were then incubated for 1.5 h. Afterwards, the cells were washed with PBS (1 x 2 mL) and fixed with paraformaldehyde (4% in PBS) for 10 min at RT. The samples were then washed with PBS (3 x 2 mL) and permeabilized with 0.1% Triton X-100 for 15 min at RT. The samples were washed with PBS (3 x 2 mL), and the coverslips were mounted with ProLong Diamond with DAPI overnight before imaging.

**Th17 cell differentiation and FACS analysis.** Naïve $CD4^+$ T cells were harvested from the spleens of mice using the isolation kit according to the manufacturer's protocol. Forty eight-well plates were coated with 5 µg/mL anti-CD3 and incubated for 3 h at 37 °C. Cells were then washed with PBS (0.5 mL) once before plating the cells at $0.5\times10^6$ cells/well in 0.5 mL of RPMI-1640 media (with 10% FBS, L-glutamine, sodium pyruvate, HEPES buffer, and penicillin/streptomycin). Neutralizing antibodies (anti-IL4 and anti-IFN-γ, 10 µg/mL), anti-CD28 (2 µg/mL), and cytokines (mIL-6, 20 ng/mL; hTGF-β, 5 ng/mL) were added to induce Th17 cell differentiation (Th17 cell conditions). Cultures were treated with FICZ (300 nM), PIM-Th17 (10 μM), or vehicle (DMSO) with or without UV light (365 nm, 1 min) and incubated for 4 d at 37 °C before FACS analysis.

*FACS staining.* Cells were collected in microcentrifuge tubes and centrifuged at 350 ×g for 3 min at 4 °C. Cells were resuspended in 100 μL FACS buffer, transferred to a 96-well V bottom plate, and centrifuged at 350 ×g for 3 min at 4 °C. The cells were resuspended in 50 μL of FACS buffer containing anti-mouse CD16/CD32 (1:300) and incubated for 10 min on ice. Cells were then centrifuged at 350 ×g for 3 min at 4 °C and resuspended in 50 μL of FACS buffer containing APC anti-mouse CD4 (1:300). The cells were incubated on ice for 15 min in the dark, after which the cells were washed twice with 170 μL of FACS buffer. The cells were then fixed and permeabilized according to the manufacturer's protocol (Affymetrix eBioscience IC Fixation and permeabilization buffer set). The cells were centrifuged at 350 ×g for 4 min, resuspended in 50 μL of permeabilization buffer containing PE anti-mouse/rat IL-17A (1:100), and incubated for 30 min at RT. The cells were washed twice with 170 μL of permeabilization buffer. After the last wash, cells were resuspended in 170 μL of FACS buffer and transferred to FACS tubes for analysis.

**Statistics.** Each experiment was carried out at least three independent times. Error bars represent standard deviation from the mean. For box plots, interquartile ranges (IQRs, boxes), median values (line within box), whiskers (lowest and highest values within 1.5 times IQR from the first and third quartiles), and outliers beyond whiskers (dots), are shown. Statistical significance was assessed using one-way ANOVA followed by post-hoc Tukey's test.

**Table S1. Primer sequences for qPCR**

| Gene | Forward primer | Reverse primer |
|---|---|---|
| *hCyp1a1* | GTGATCCCAGGCTCCAAGAG | AGAAGAAACTCCGTGGCCG |
| *hCyp1a2* | GCTGAATGGCTTCTACATCCCC | GCGGTGAGGAACCGCTC |
| *hCyp1b1* | AACGTACCGGCCACTATCAC | GCACTCGAGTCTGCACATCA |
| *hAhrr* | GCAGCGGAGATGAAAATGAGG | TTCCGATTCGCACAGACTGG |
| *hRpl13a* | TCCTCCTTTTCCAAGCGGC | GGCCCAGCAGTACCTGTTT |

**Synthetic procedures**

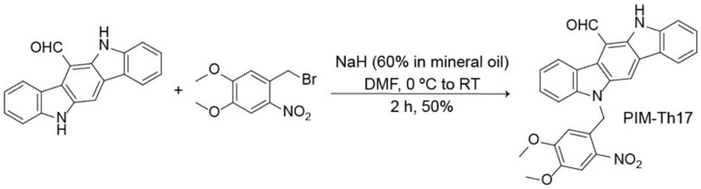


**Scheme S1. Synthesis of PIM-Th17**

**PIM-Th17** (**P**hoto-activatable **I**mmune **M**odulator of **Th17** cells). To 10 mg of FICZ[4–6] (0.035 mmol, 1.0 eq.) was added 350 μL of anhydrous DMF, and the solution was cooled down to 0 °C. The mixture was stirred for 5 min prior to adding 60% NaH (2.1 mg, 0.053 mmol, 1.5 eq.) and stirred for an additional 10 min. To this mixture was added NV-Br (12 mg, 0.043 mmol, 1.2 eq.), and the reaction was stirred for 2 h at RT. The reaction was quenched by adding 2 mL of $ddH_2O$ at 0 °C. Fifty milliliters of $ddH_2O$ was added to the mixture, and the organic layer was extracted with 10 mL of ethyl acetate twice. The combined organic layers were washed with $ddH_2O$, dried over anhydrous $Na_2SO_4$, and concentrated under reduced pressure. The resulting crude product was purified by silica gel flash chromatography, eluting with 80:20 hexane:ethyl acetate to yield a bright yellow solid, which was further purified by HPLC to obtain 8.5 mg of PIM-Th17 (50% yield). $^1$H NMR (600 MHz, DMSO-d6): δ 11.83 (s, 1H), 11.43 (s, 1H), 8.87 (s, 1H), 8.69 (d, 1H, J = 12 Hz), 8.21 (d, 1H, J = 12 Hz), 7.86 (s, 1H), 7.75 (d, 1H, J = 12 Hz), 7.63 (d, 1H, J = 12 Hz), 7.52 (t, 1H, J = 8 Hz), 7.44 (t, 1H, J = 8 Hz), 7.30 (t, 1H, J = 8 Hz), 7.21 (t, 1H, J = 8 Hz), 6.21 (s, 1H), 5.72 (s, 1H), 3.85 (s, 3H), 3.03 (s, 3H). $^{13}$C NMR (500 MHz, DMSO-d6): δ 190.67, 147.94, 142.15, 140.22, 129.03, 127.48, 126.93, 125.42, 120.92, 120.29, 119.81, 112.66, 110.51, 109.33, 109.23, 109.05, 56.59, 55.90. ESI-MS: Calcd. For $C_{28}H_{22}N_3O_5^+$ $(M+H)^+$ 480.1559, found 480.1563.

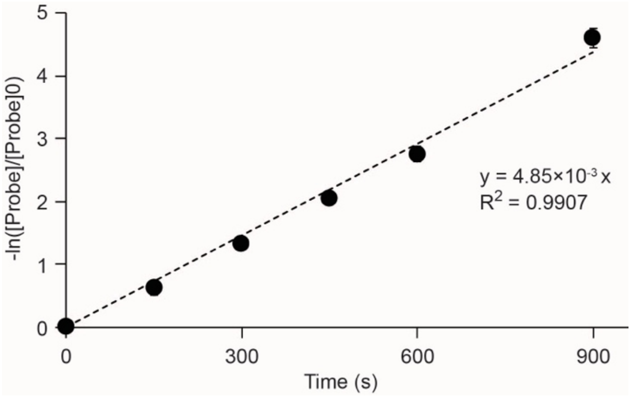


**Figure S1. Photo-uncaging of PIM-Th17 is a first-order reaction.** PIM-Th17 was irradiated with 365 nm UV light for 15 min. The reaction was periodically analyzed by HPLC and MS to monitor conversion of PIM-Th17 to FICZ. Percent conversion was calculated by analyzing standards of PIM-Th17 and FICZ by HPLC, using integration of the area under the curve for normalization. Error bars represent mean ± SD.

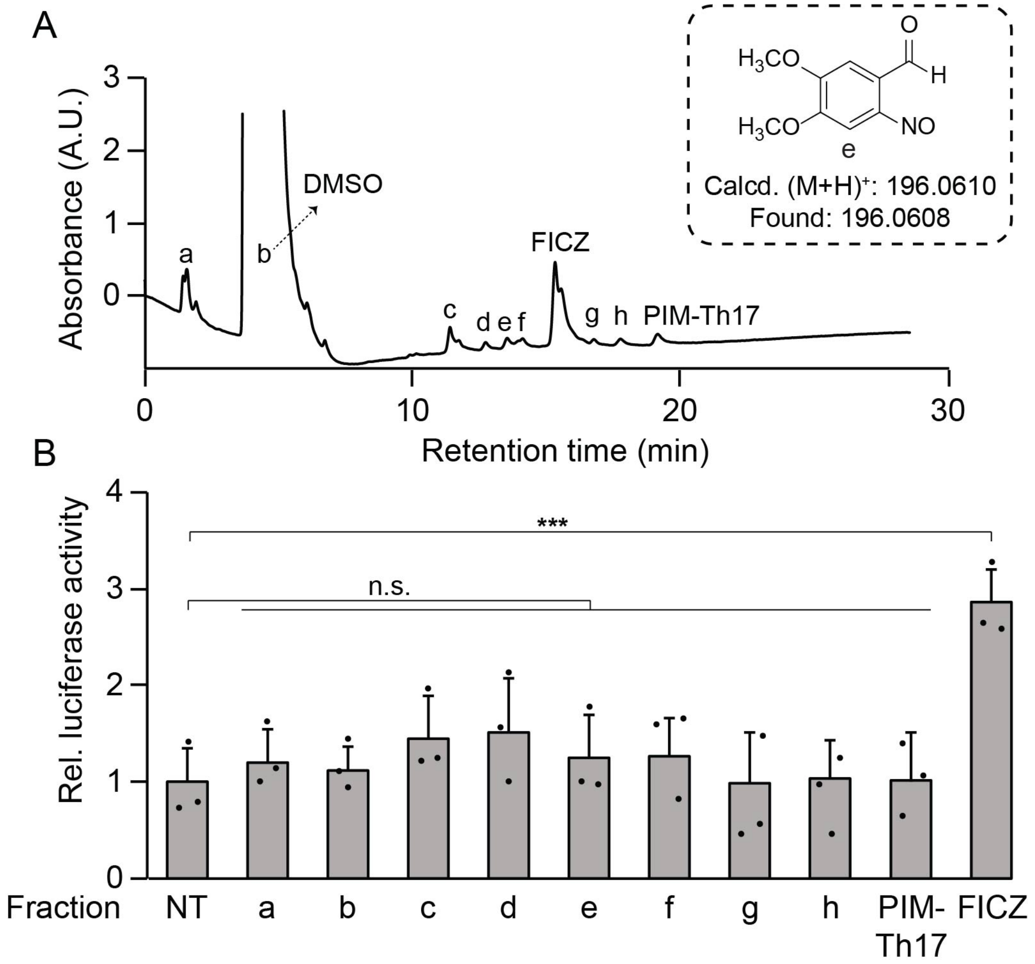


**Figure S2. Photo-uncaging of PIM-Th17 produces minor byproducts that do not activate AhR.** (A) PIM-Th17 (10 µM) was photo-uncaged in vitro, and the byproducts were purified by HPLC and analyzed by MS. (B) AhR luciferase reporter cells (HEK 293T cells co-transfected with pGudluc6.1 and pRLTK) were treated with or without the isolated byproducts for 2 h, and the luminescence was measured. Data are representative of three independent experiments. Error bars represent mean ± SD. n.s. = not significant, ***p<0.001

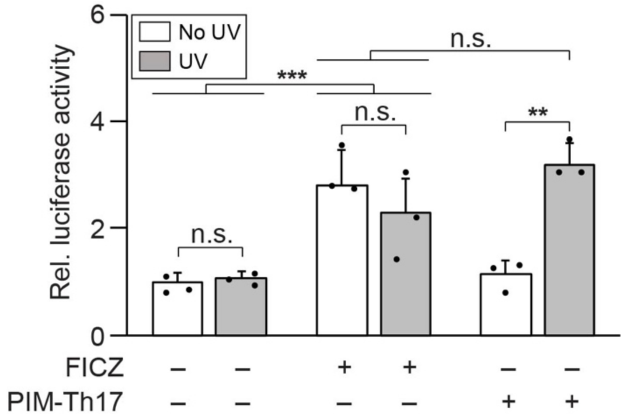


**Figure S3. PIM-Th17 does not increase AhR activity until photo-uncaged.** PIM-Th17 (10 μM), FICZ (300 nM), or vehicle (DMSO) were treated with and without UV light (365 nm, 10 min). HEK 293T cells expressing an AhR luciferase reporter (Figure S2) were treated with these samples, and the resulting luminescence was measured after 2 h. Data are representative of three independent experiments. Error bars represent mean ± SD. n.s. = not significant, **p<0.01, ***p<0.001.

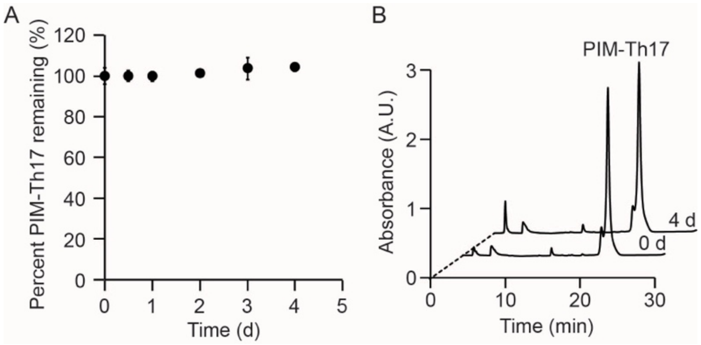


**Figure S4. PIM-Th17 is stable in aqueous solution.** PIM-Th17 was dissolved in a mixture of 5:1 DMSO:PBS and incubated at 37 °C for 4 d. (A) Percent remaining of PIM-Th17. (B) HPLC analyses of samples. Traces shown were monitored by UV absorbance at 254 nm. A.U. = arbitrary units. Data are representative of three independent experiments. Error bars represent mean ± SD.

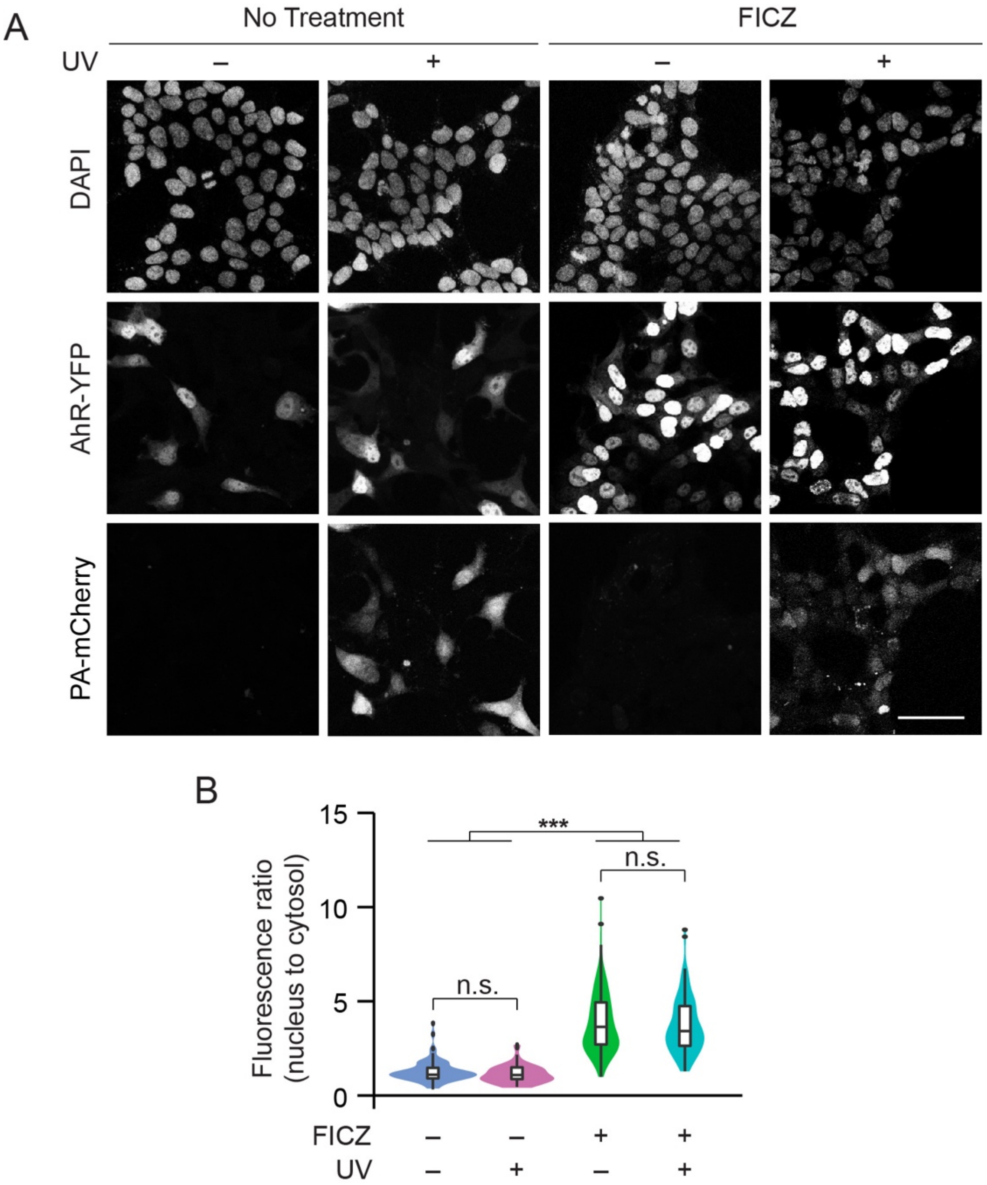


**Figure S5. FICZ leads to nuclear translocation of AhR-YFP.** (A) HEK 293T cells co-transfected with AhR-YFP and PA-mCherry were pre-treated with FICZ (300 nM) or vehicle (DMSO) for 30 min, and the center of the dish was irradiated with UV light (epifluorescence using DAPI filter, 30 s). Cells were subsequently incubated for 1.5 h, fixed, stained with DAPI, and imaged. (A) Maximum intensity z-projection images of cells at the periphery (- UV) or center (+ UV) of the dish, imaged by confocal microscopy. Scale bar: 50 μm. (B) Violin plot of nuclear to cytosolic ratio of AhR-YFP from (A), n = 50–70 cells in each treatment group. n.s. = not significant, ***$p<0.001$.

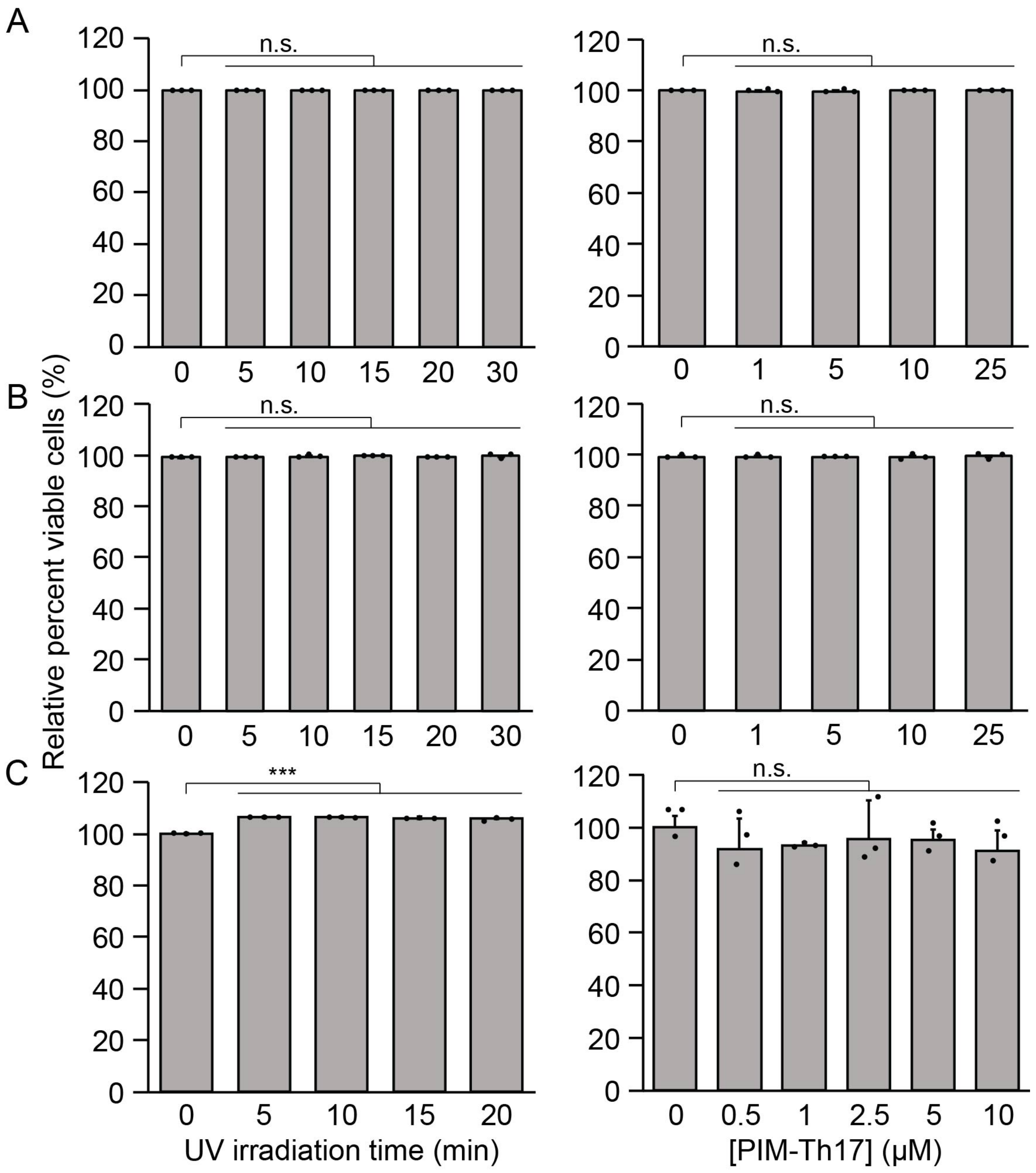


**Figure S6. Photo-uncaging conditions are not toxic to cells.** Cell viability was measured by propidium iodide staining and FACS analysis. (A) HEK 293T, (B) HepG2, and (C) mouse splenic CD4$^+$ T cells were exposed to UV light (365 nm) for various amounts of time (left) or different concentrations of PIM-Th17 (A-B) for 24 h or (C) 4 d (right). Data are representative of three independent experiments. Error bars represent mean ± SD. n.s. = not significant, ***$p<0.001$.

¹H NMR of PIM-Th17 in $(CD_3)_2SO$ (600 mHz)

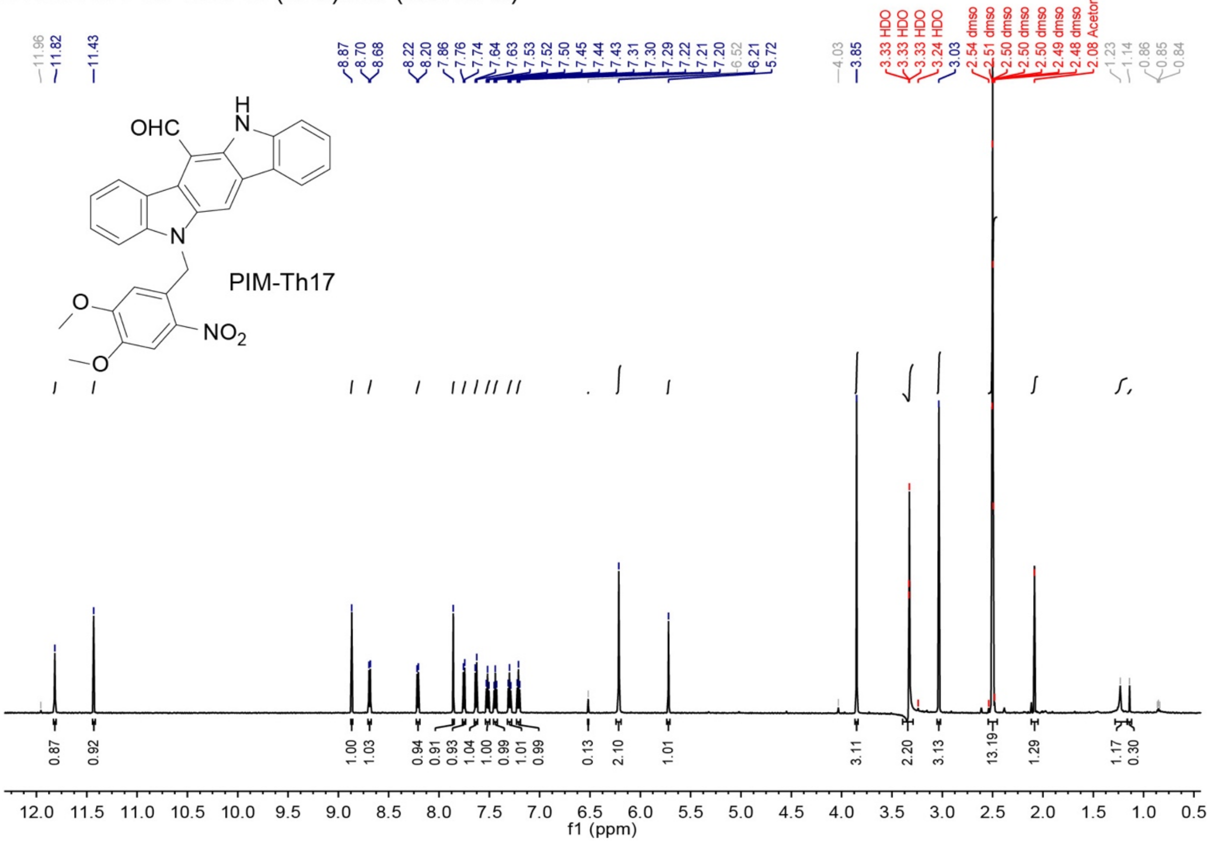


**Figure S7. $^1$H NMR spectrum of PIM-Th17**

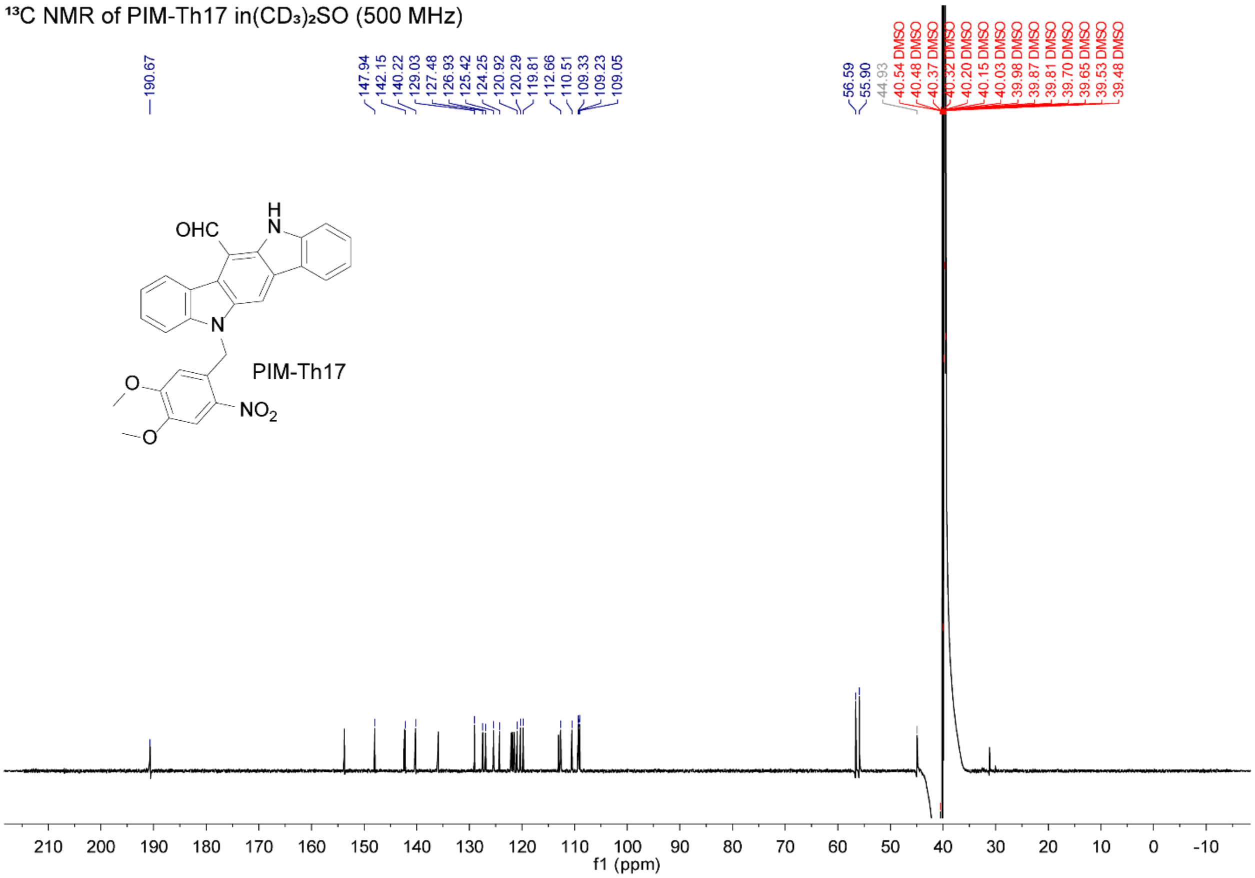


**Figure S8. $^{13}$C NMR spectrum of PIM-Th17**

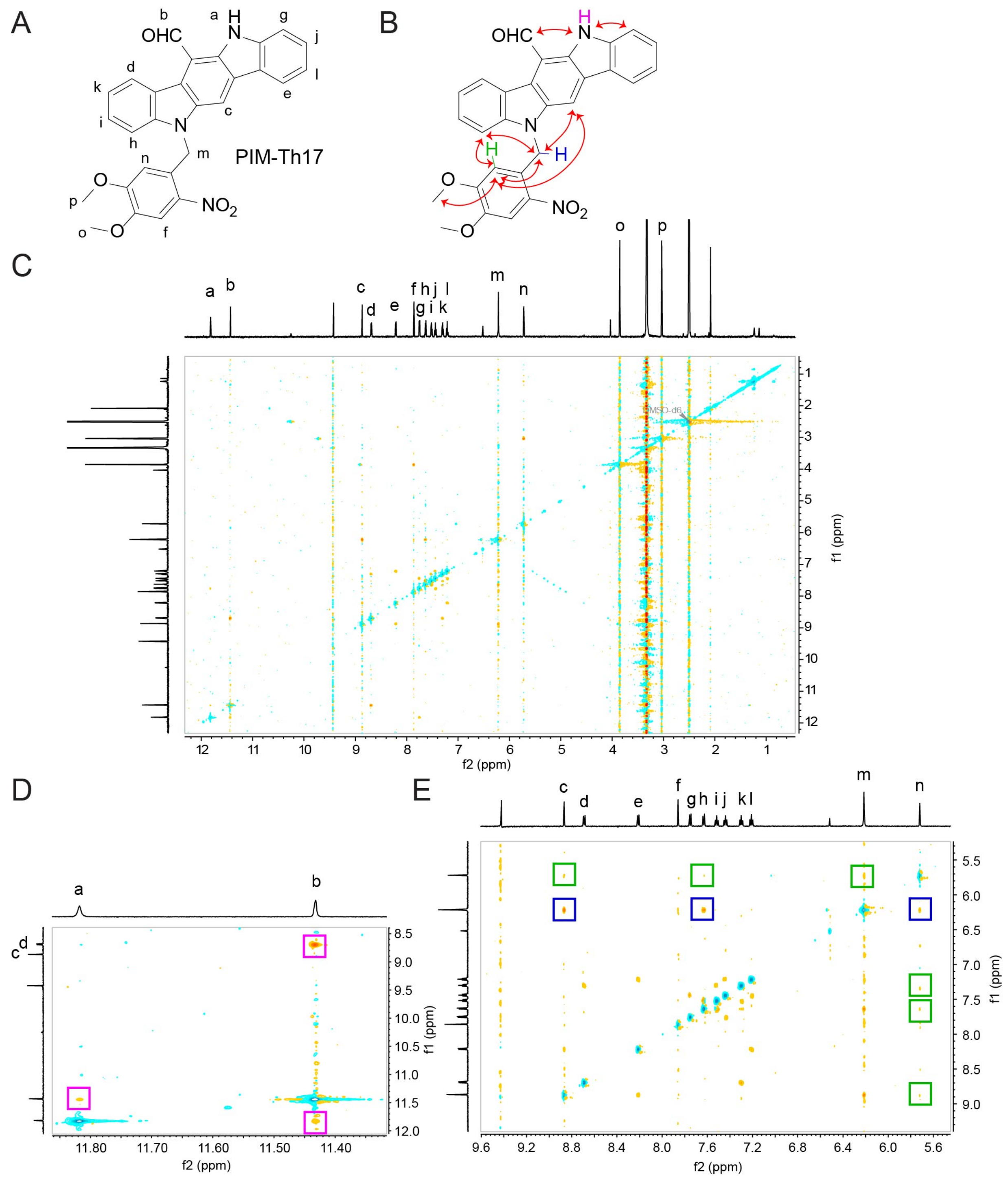


**Figure S9. 2D ROESY spectrum of PIM-Th17.** (A) Structure of PIM-Th17 with $^{1}$H assignments indicated. (B) Important $^{1}$H correlations for structure determination. (C) 2D ROESY spectrum. (D) Enlarged regions of 2D ROESY spectrum to highlight important $^{1}$H correlations.
